# Predicting the impacts of sea level rise in sea turtle nesting habitat on Bioko Island, Equatorial Guinea

**DOI:** 10.1101/760538

**Authors:** Callie A. Veelenturf, Elizabeth M. Sinclair, Frank V. Paladino, Shaya Honarvar

## Abstract

Sea level is expected to rise 44 to 74 cm by the year 2100, which may have critical, previously un-investigated implications for sea turtle nesting habitat on Bioko Island, Equatorial Guinea. This study investigates how nesting habitat will likely be lost and altered with various increases in sea level, using global sea level rise (SLR) predictions from the Intergovernmental Panel on Climate Change. Beach profiling datasets from Bioko’s five southern nesting beaches were used in GIS to create models to estimate habitat loss with predicted increases in sea level by years 2046-2065 and 2081-2100. The models indicate that an average of 62% of Bioko’s current nesting habitat could be lost by 2046-2065 and 87% by the years 2081-2100. Beach D is predicted to be the least vulnerable to increases in sea level. Erosion and tall vegetation berms have been documented on Beaches A and B, causing green turtles to nest uncharacteristically in front of the vegetation line. Development plans are currently underway for Beach D. With Beach D being the least susceptible to future increases in sea level, development and anthropogenic encroachment here would be especially detrimental to nesting turtle populations. Identified habitat sensitivities to SLR will be used to inform the government of Equatorial Guinea to consider the vulnerability of their resident turtle populations and projected climate change implications when planning for future development. To our knowledge this is the first study to predict the impacts of SLR on sea turtle nesting habitat in Africa.

## Introduction

One of the important discussions involving climate change and sea turtle conservation is the imminent loss of sea turtle nesting habitat corresponding to increasing sea level rise (SLR) [1, 2, 3, 4]. The Intergovernmental Panel on Climate Change (IPCC) has generated four scenarios that predict SLR for years 2046-2065 and 2081-2100. The IPCC indicates that an increase of about 44 to 74 cm will be experienced globally by the year 2100, which will have previously un-investigated implications for the second largest nesting aggregations of leatherback (*Dermochelys coriacea*) and green sea turtles (*Chelonia mydas*) in West Africa [5–9]. Small islands are at the greatest risk from climate change [10, 11], and expected effects include increased salinity within the water table, beach erosion, and sand inundation with increased tide elevation [12, 13]. Marine reptiles have evolved with natural coastal erosive processes such as high-tide flooding, accretion, and seasonal erosion, but the extreme beach modification of the past half-century is progressing at an alarming rate, faster than the sexual maturation rate of some species [1]. Great Barrier Reef green sea turtle nesting population, the largest in the world, has experienced hatchling success reduction in the past few years, which is thought to be a result in part of a rising groundwater table [14]. It has been estimated that the most extreme SLR predictions will result in inundation of 27 percent of Great Barrier Reef green sea turtle nesting habitat [3]. Nesting habitat inundation is also expected in Bonaire (26%) and Barbados (32%) with a 0.5 m rise in sea level [1, 2]. A study conducted on the Nile Delta using GIS modeling and IPCC predictions suggest a maximum of 49.22% of the coast is susceptible to inundation [4].

For nesting habitat that is not inundated, SLR will undoubtedly alter the potential for previous nesting beaches to continue to maintain their historic [1]. With an overall reduction in nesting habitat, the density of nests will likely increase within the area of available nesting habitat. This has potential to cause decreased hatching success through increased contamination and physical disturbance of nests by co-specifics [2, 14–17]. Since species can shift their nesting grounds when faced with unsuitable nesting habitats [18], it is important to also investigate multiple nesting areas within a single species in a nesting region [3]. It has been suggested that with median expected SLR in the Hawaiian Islands, green sea turtle nesting localities will likely need to shift primarily from Trig, Gin, and Little Gin islands to East Island in order for historic reproductive productivity to be sustained [19]. With increasing SLR, increases in erosion rates and nests that are flooded from storms can be expected [11, 20, 21]. Effects from an increased water table due to SLR can already be observed on Raine Island, Australia, where depressions from sea turtle body pits have been observed filling with water [18, 22]. This increased nest inundation will undoubtedly cause decreases in reproductive output of all sea turtle species [23, 24].

The 10.75 km of main sea turtle nesting beaches (A-E, Fig 1) on the southern side of Bioko Island are critically important nesting habitat for the leatherback and green sea turtles in the West/Central African region [9, 25–27]. Green turtles nest mostly on beaches A, B and C, and leatherback sea turtles on C, D and E [9]. Within and among species, there is variation in selection for more specific beach characteristics such as beach length, width, height, slope, orientation, and vegetation [28–31]. The various beach types where sea turtles nest combined with the specific nest-site characteristics that are selected for by each species can be altered in diverse ways by increasing sea level [1]. Green turtles prefer to nest on narrower, steeper beaches and in the area behind the vegetation line, whereas leatherback sea turtles prefer wider, flatter beaches and the area between the high tide line (HTL) and the vegetation line. It has been found previously that narrower beaches at lower elevations are more susceptible to SLR [1]. As the morphology of the beaches and intricate beach zoning is altered, these habitat selection differences cause species-specific SLR threats. Based on the spatial distribution of nests within each species, this threat could be more severe and more imminent for some species than others.

**Fig 1.**
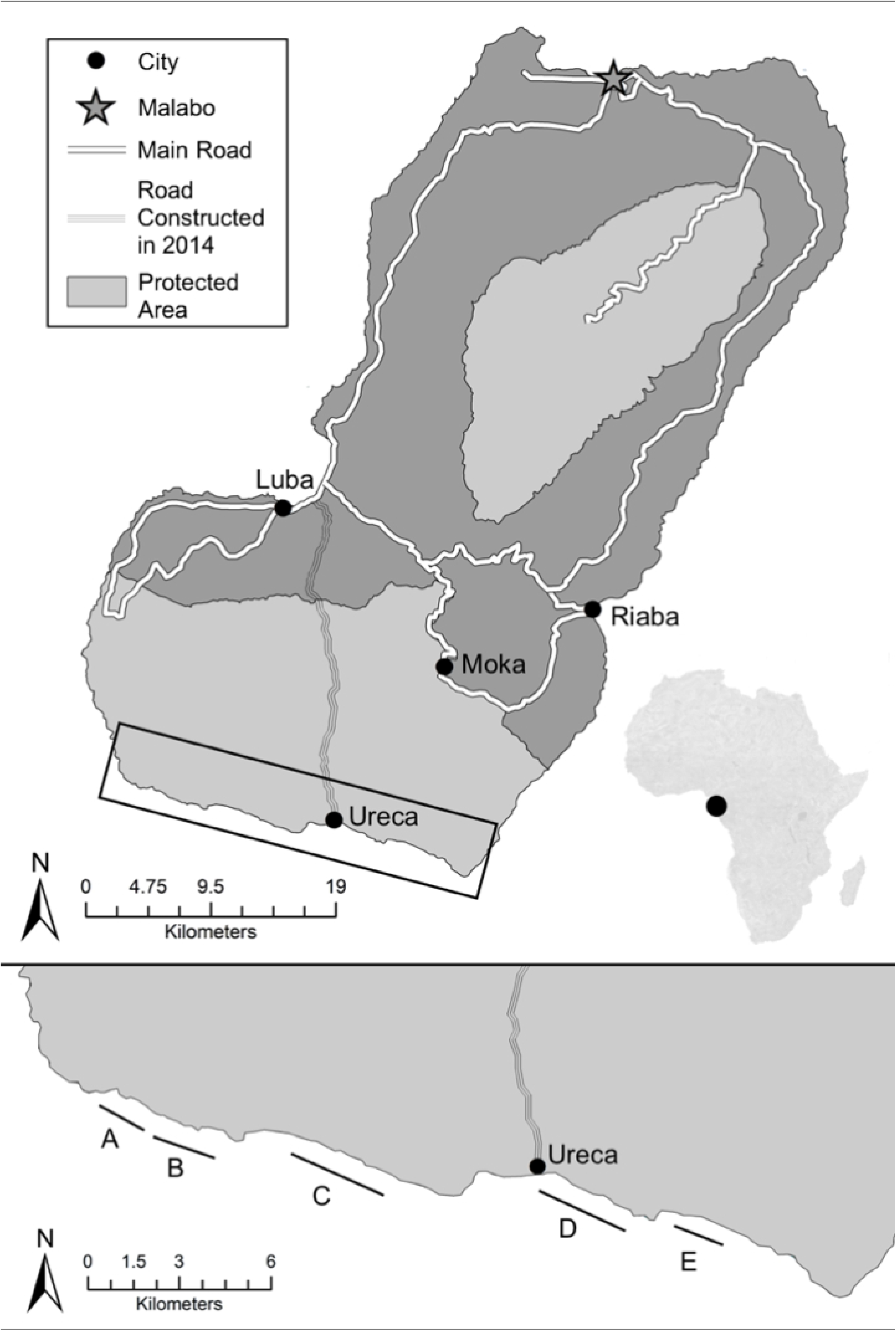
Bioko Island Nesting Beaches. The five nesting beaches are labelled with letters A–E.

The goal of this study is to characterize sea turtle nesting beaches on Bioko Island and to model the effects of SLR for use in generating targeted conservation management plans. Nest locations from both green and leatherback turtles were used together with SLR predictions on Bioko’s 5 nesting beaches to determine how each species will likely be affected in the upcoming years by climate change. Our objectives were to (1) construct a 3D profile of 5 nesting beaches by collecting morphometric/contour data in an x, y, z dimensional space, (2) use triangulated irregular network (TIN) models and digital elevation models (DEM) to map landward movement of the HTL, and (3) predict how the model output will affect green and leatherback turtle nesting on Bioko Island.

## Materials and methods

### Data collection

Beach profiling transects were conducted on all five of Bioko’s nesting beaches (A-E) (Fig 1, Tables 1–3). Beach characterization methods were consistent with a similar SLR prediction model for 13 beaches on the island of Bonaire, Dutch Antilles [1]. The profile of each beach was recorded relative to the high-water mark at 50 m intervals along the beaches using a 60 m measuring tape. The transects on each beach were 50 m apart, perpendicular to the water line, and spanned the distance from the vegetation line to the drop off during lowest tide. A clinometer, meter tape, and compass were used to create profiles of beach topography and dimensions at each change in slope along the transect. Accuracy to ground truth was relative to the stake GPS point (Garmin GPSMap 64) at the start of each transect. To insure maximum accuracy, up to 6 different waypoints for the same stake on each beach were averaged to generate an average stake reference point to be used in the following spatial analysis. Beach D was profiled once at the beginning of the season and once at the end to better understand how seasonal fluctuations could affect SLR predictions for a single beach. The circular error probable for each stake was calculated. This work was conducted under appropriate permits from Universidad Nacional de Guinea Ecuatorial (#289/2016) and the Institutional Animal Care and Use Committee at Purdue University (IACUC protocol #1410001142).

**Table 1.** Official Beach Locations.

| BEACH | START |  | END |  |
| --- | --- | --- | --- | --- |
|  | Latitude | Longitude | Latitude | Longitude |
| **A** | N 3 16.248 | E 8 27.610 | N 3 15.850 | E 8 28.456 |
| **B** | N 3 15.674 | E 8 28.790 | N 3 15.357 | E 8 29.842 |
| **C** | N 3 15.197 | E 8 31.600 | N 3 14.517 | E 8 33.321 |
| **D** | N 3 14.762 | E 8 35.550 | N 3 14.092 | E 8 36.871 |
| **E** | N 3 14.032 | E 8 37.872 | N 3 13.726 | E 8 38.697 |

GPS points for the start and end of beaches A-E, from Northwest to Southeast.

**Table 2.** Beach Morphometrics 1.

| Beach | Length (km) | Total Area (m<sup>2</sup>) | Current Nesting Habitat (m<sup>2</sup>)<br>(Proportion of Total Beach Area) |
| --- | --- | --- | --- |
| A | 1.7 | 84,914 | 8,852 (0.10) |
| B | 1.9 | 145,350 | 10,350 (0.07) |
| C | 2.9 | 236,784 | 23,946 (0.10) |
| D | 2.65 | 350,564 | 63,592 (0.18) |
| E | 1.6 | 153,110 | 16,217 (0.11) |

Morphometrics of Bioko's five nesting beaches. Nesting habitat is defined as the area between the high tide line and vegetation line.

**Table 3.** Beach Morphotmetrics 2.

| Beach | <u>Max.</u><br><u>Elevation</u><br><u>(m)</u> | <u>Min.</u><br><u>Elevation</u><br><u>(m)</u> | <u>Average</u><br><u>Elevation</u><br><u>(m) ±SD</u> | <u>Max</u><br><u>Width</u><br><u>(m)</u> | <u>Min</u><br><u>Width</u><br><u>(m)</u> | <u>Mean</u><br><u>Width</u><br><u>(m) ±SD</u> |
| --- | --- | --- | --- | --- | --- | --- |
| A | 1.62 | 1.13 | 0.37 ± 0.53 | 100.23 | 16.9 | 49.06 ± 14.87 |
| B | 1.85 | -1.01 | 0.57 ± 0.48 | 158.24 | 9.68 | 70.67 ± 31.03 |
| C | 1.80 | -1.03 | 0.51 ± 0.70 | 137.14 | 29.44 | 78.80 ± 23.65 |
| D | 1.77 | -2.08 | 0.57 ± 0.71 | 215.49 | 81.3 | 126.076 ± 31.70 |
| E | 1.30 | -0.18 | 0.78 ± 0.36 | 154.65 | 59.93 | 94.80 ± 18.15 |

Morphometrics of Bioko's 5 nesting beaches based on 2017 profiling data. Averages show ± standard deviation.

### Spatial analysis

A program was written using Python to perform calculations on beach profiling data to generate a waypoint and elevation at each change in slope on the transects and at the present elevation of the high tide line. In ArcMap (Esri version 10.4), GPS points with their respective elevation values were entered as x, y, z data and then projected as shapefiles. All elevations were relative to the HTL, which for the purposes of this project is at an elevation of 0 m. The points to line feature in ArcMap was used to create 5 lines connecting: 1) stake GPS points, 2) HTL GPS points, 3) GPS points of the final segment of each transect, 4) GPS points for transect 1 on each beach, and 5) all points on the last transect for each beach. These lines allowed the “feature to polygon” tool to be used to create a polygon of each beach. The waypoints from all changes in slope on all transects were then used as inputs to the topo to raster tool to create a DEM. The raster dataset was then used in the creation of a TIN, for 3D visualizations of beach morphology and changes due to SLR [1, 3]. The raster datasets were projected and reclassified to reflect the IPCC predictions of climate change (0.24, 0.25, 0.26, 0.30, 0.4, 0.48, 0.63 and 0.75 m) [8] (Tables 4 and 5). The range of one class of each raster always ended at 0 m, so the current approximate viable nesting habitat could be easily isolated. For the purpose of this study, nesting habitat was defined by beach area between the high tide line and vegetation line. The extract by mask feature was then used to clip these rasters to the polygons of each beach. The count of each class along with the cell size was used to calculate the area of each beach, area of current nesting habitat, and area lost and left under each SLR predation. ArcScene (Esri version 10.4) was used to create 3D graphics of models, and ArcGIS software was used to generate all maps and basemaps. Species-specific predictions of impacts of climate change were then made based on the spatial presence of each species on each beach coupled with the beach’s vulnerability to SLR.

**Table 4.** Habitat Loss 2046-2065.

| **Nesting Habitat Inundated (proportion of total nesting habitat)** |  |  |  |  |  |
| --- | --- | --- | --- | --- | --- |
| **Beach** | **0.24 m** | **0.25 m** | **0.26 m** | **0.30 m** | **Mean** |
| **A** | 6,209 (0.70) | 6,344 (0.72) | 6,444 (0.73) | 6,835 (0.77) | 0.73 |
| **B** | 8,507 (0.78) | 8,633 (0.79) | 8,760 (0.80) | 9,282 (0.85) | 0.81 |
| **C** | 13,422 (0.50) | 13,851 (0.52) | 14,245 (0.53) | 15,802 (0.59) | 0.54 |
| **D** | 29,246 (0.45) | 30,092 (0.46) | 30,938 (0.48) | 34,522 (0.53) | 0.48 |
| **E** | 9,404 (0.50) | 9,807 (0.52) | 10,151 (0.53) | 11,373 (0.60) | 0.54 |

The potential area (m<sup>2</sup>) on 5 of Bioko's nesting beaches that would be lost to sea level rise (SLR) under 4 scenarios for 2046-2065: 0.24, 0.25, 0.26, and 0.30 m. The average represents an average SLR loss predicted by the 4 scenarios for 2046-2065.

**Table 5.** Habitat Loss 2081-2100.

| **Nesting Habitat Inundated (proportion of total nesting habitat)** |  |  |  |  |  |
| --- | --- | --- | --- | --- | --- |
| **Beach** | **0.4 m** | **0.47 m** | **0.48 m** | **0.63 m** | **0.75 m** |
| **A** | 7,544 (0.85) | 7,851 (0.89) | 7,887 (0.89) | 8,239 (0.93) | 8,396 (0.95) |
| **B** | 9,576 (0.93) | 9,960 (0.96) | 9,996 (0.97) | 10,296 (0.99) | 10,338 (1) |
| **C** | 18,282 (0.76) | 20,052 (0.84) | 20,280 (0.85) | 22,494 (0.94) | 23,328 (0.97) |
| **D** | 42,782 (0.67) | 47,314 (0.74) | 47,885 (0.75) | 54,761 (0.86) | 58,542 (0.92) |
| **E** | 12,761 (0.79) | 13,723 (0.85) | 13,722 (0.85) | 15,313 (0.94) | 15,869 (0.98) |

The potential area (m<sup>2</sup>) on 5 of Bioko's nesting beaches that would be lost to sea level rise (SLR) under 4 scenarios for 2081-2100: 0.4, 0.47, 0.48, and 0.63, and 1 scenario for 2100, 0.75 m. The mean represents an average SLR loss predicted by the 4 scenarios for 2081-2100.

## Results

The average circular error probable for the reference points was 2.43 ± 1.44 (standard deviation), indicating that 2.43 m is the radius of a circle centered around the mean position of each reference stake that contains 50% of the reference stake GPS points. Similarly, the circular error probable was 3.85 ± 2.51 for 98% of the reference stake GPS points. Satellite views of the transects confirmed their proper spacing and alignment.

Beach A is the smallest beach with an average beach width of 49.06 m and a total beach area of 84,914 m^2^ (Tables 2 and 3). Beach A also had the smallest area of nesting habitat in 2017, 8,852 m^2^, but Beach B has the smallest percentage of nesting habitat to total beach area, 7% (Table 2). Beaches C and D were the two longest and widest nesting beaches on Bioko Island, with areas of 236,784 m^2^ and 350,564 m^2^, respectively (Table 2). Beach D contained the highest percentage of nesting habitat out of all 5 beaches, 18% (Table 2). This beach was also the closest to the road built in 2014 and is thus one of the beaches that is the most vulnerable to illegal egg and adult turtle take and development (Fig 1). Satellite imagery and photographs of Beaches D showed the evident discrepancy across nesting beaches in terms of nesting habitat available in 2017. The average elevation of Beach A (farthest west) was the lowest (0.37 m) and that of Beach E (farthest east) the highest (0.77 m) (Table 3). Beach A was the steepest beach, and Beach D was the shallowest, as expected (Fig 2). Beach A was the only beach that had a significantly different slope than all other beaches (Fig 2).

**Fig 2.**
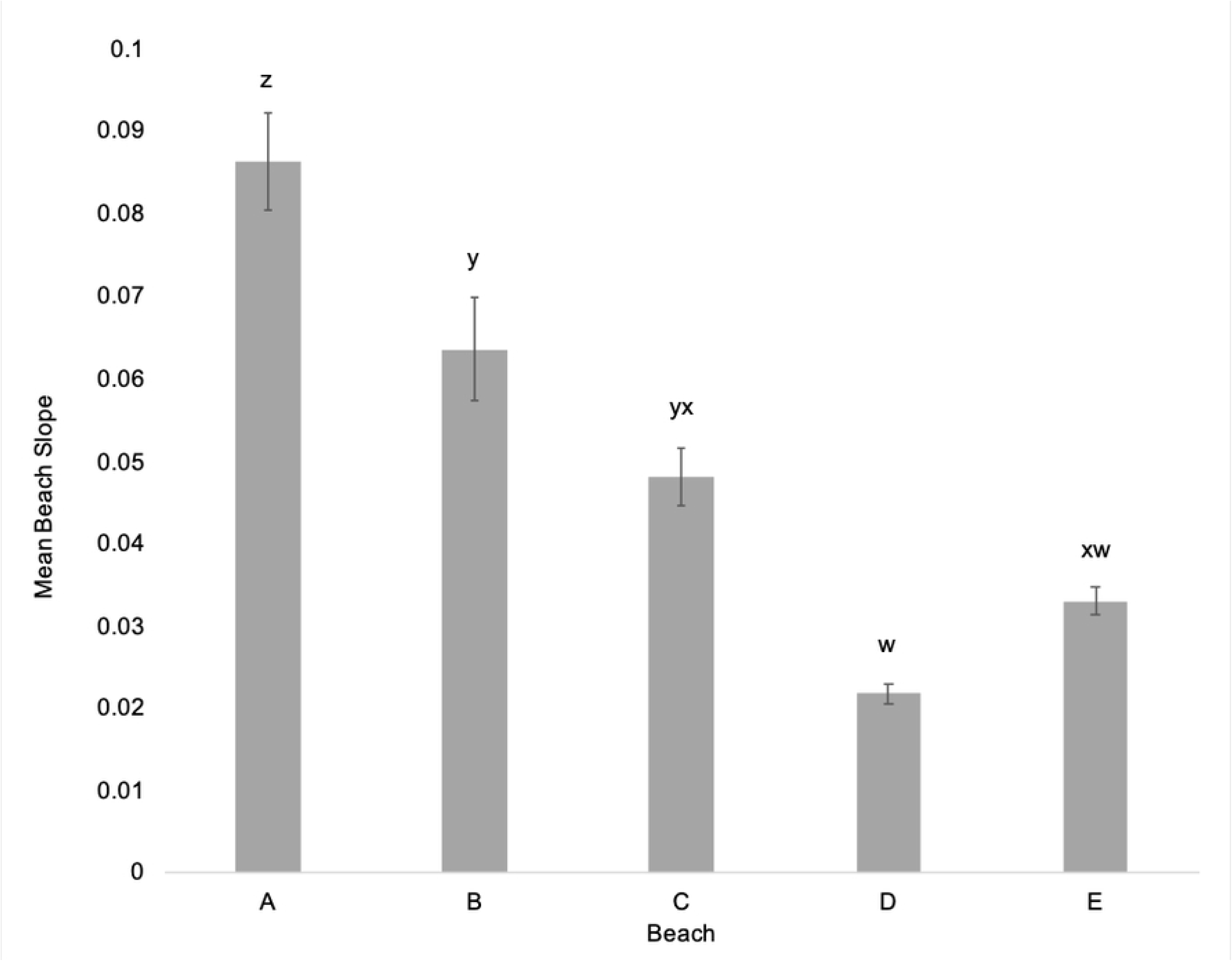
Mean Beach Slopes. The slope for each transect was measured from the vegetation line to the drop off at lowest tide and slopes for all transects were averaged per beach. Differing letters indicate significant differences between mean beach slopes. Error bars are standard error from the means.

There are four scenarios from the Intergovernmental Panel on Climate Change (IPCC) that predict SLR for years 2081-2100 and for the years 2046-2065. The results presented here are calculated with average SLR under Representative Concentration Pathways (RCP) 2.6, 4.5, 6.0, and 8.5 for 2081-2100, the average SLR under RCP2.6, 4.5, 6.0, and 8.5 predictions for 2046–2065, and the average SLR under the RCP8.5 prediction specifically for 2100. The RCP8.5 prediction for 2100 was included to show the most extreme extent of current IPCC predictions.

Within only 30 years, using different scenarios of SLR, these models predict drastic changes in beach morphology. Under the most extreme scenario for 2046-2065, with a 0.3 m increase in sea level, Beach D is predicted to lose the least amount of its current nesting habitat, only 53%, and Beach B is expected to lose the most with a predicted 81% nesting habitat loss (Table 4). Under the least extreme scenario, all beaches will lose at least 45% of its current nesting habitat, and Beach B is likely to lose 78% of its current available nesting habitat (Table 4). The beaches where green sea turtles nest in greater quantities, A and B, will experience higher nesting habitat losses than those where leatherback sea turtles nest more often, D and E (Table 4). Under the least severe scenario, the largest proportion of current nesting habitat that would be left by 2046-2065 was 55% on Beach D, and the smallest proportion of nesting habitat that would remain on Beach B is 22% (Table 5).

Under the RCP8.5 predictions for SLR in year 2100, all beaches were predicted to lose at least 92% of their current nesting habitat (Table 5). For the average SLR across RCP2.6, 4.5, 6.0 and 8.5 scenarios for 2081-2100, no beach was predicted to lose less than 67% (Table 5). Under the most extreme scenario, Beach B is predicted to be completely inundated (Table 5). The beach expected to lose the least amount of nesting habitat is Beach D with a predicted 92% loss by the year 2100 (Table 5). Beach D is the largest and widest beach but also contains the lowest minimum elevation of the south side of the island (Tables 2 and 4). The total area across all five nesting beaches that is predicted to remain on Bioko Island for nesting under the most extreme scenario is about 6,428 m^2^, which is only about 5.23% of the nesting habitat currently available. Under the least extreme scenario, approximately 31,998 m^2^ of nesting habitat is likely to be viable, 26.02% of current habitat estimates (Table 5). Beaches typically characterized as green sea turtle nesting habitat (Beaches A, B, and C) face an average of 90% nesting habitat loss for 2081-2100, and those of leatherback sea turtles (Beaches C, D and E) face an average loss of 82% (average of 0.4, 0.47, 0.48, and 0.63 m predictions).

Beach D was profiled twice within the same season, and the results show how these predictions could fluctuate. The first time the analysis was conducted with the first set of data, the maximum predicted habitat loss for Beach D was 91.84%. The second analysis, conducted with the second set of data, revealed a habitat loss of 92.06%. This is a percent difference of 0.24% and is considered negligible for this study’s specific objectives.

Narrower, more elevated, and steeper beaches appear to be more vulnerable to climate change. There was no significant relationship between maximum (F = 0.03, df = 1,3, R^2^ = 0.01, p = 0.073), minimum (F = 1.184, df = 1,3, R^2^ = 0.30, p < 0.001) or average elevation (m) (F = 0. 03, df = 1,3, R^2^ = 0.01, p = 0.013) with average nesting habitat loss (proportion of current total nesting habitat). There was a positive, significant relationship between minimum elevation of vegetation and maximum habitat loss (Fig 3). As the minimum elevation increased, the habitat loss increased as well. The data shows significant negative relationships between minimum beach width (Fig 4) and average beach width (F=4.61, R^2^ = 0.61, p=0.001) with average habitat loss (proportion of current total nesting habitat). As the beaches become wider, the average habitat loss decreases. A significant positive relationship between average slope and average habitat loss was observed (Fig 5). Beach D, the beach expected to lose the least of its current nesting habitat, has the flattest slope and the lowest minimum elevation (Fig 2, Table 3). All beaches had significantly different slopes, but Beaches A and B were the steepest, and D and E were the shallowest, as expected (Fig 2). During the highest tide during the full moon in November 2016, there was no distance between the HTL and vegetation line [32]. Green turtle nests were laid in steeper and narrower sections of the beach, whereas leatherback nests were laid in shallower and wider areas (Fig 6).

**Fig 3.**
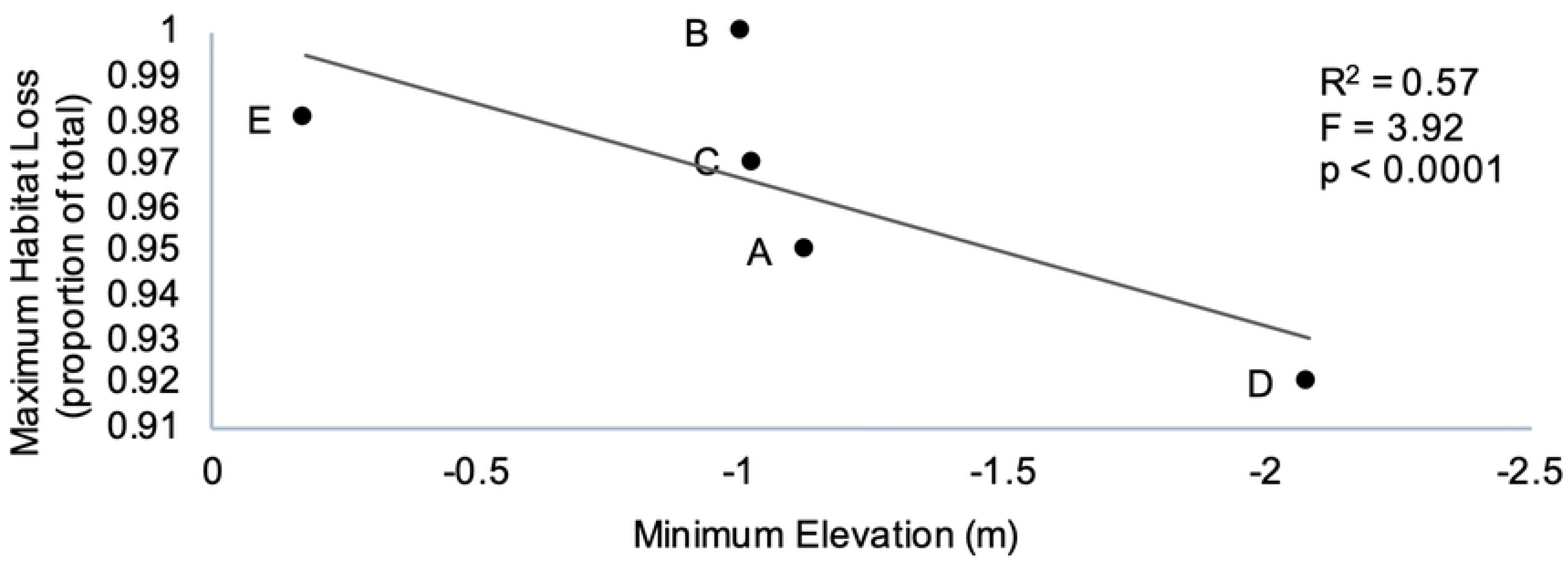
Beach Elevation versus Habitat Loss. Relationship between minimum beach elevation (m) and maximum habitat loss (proportion of total current nesting habitat).

**Fig 4.**
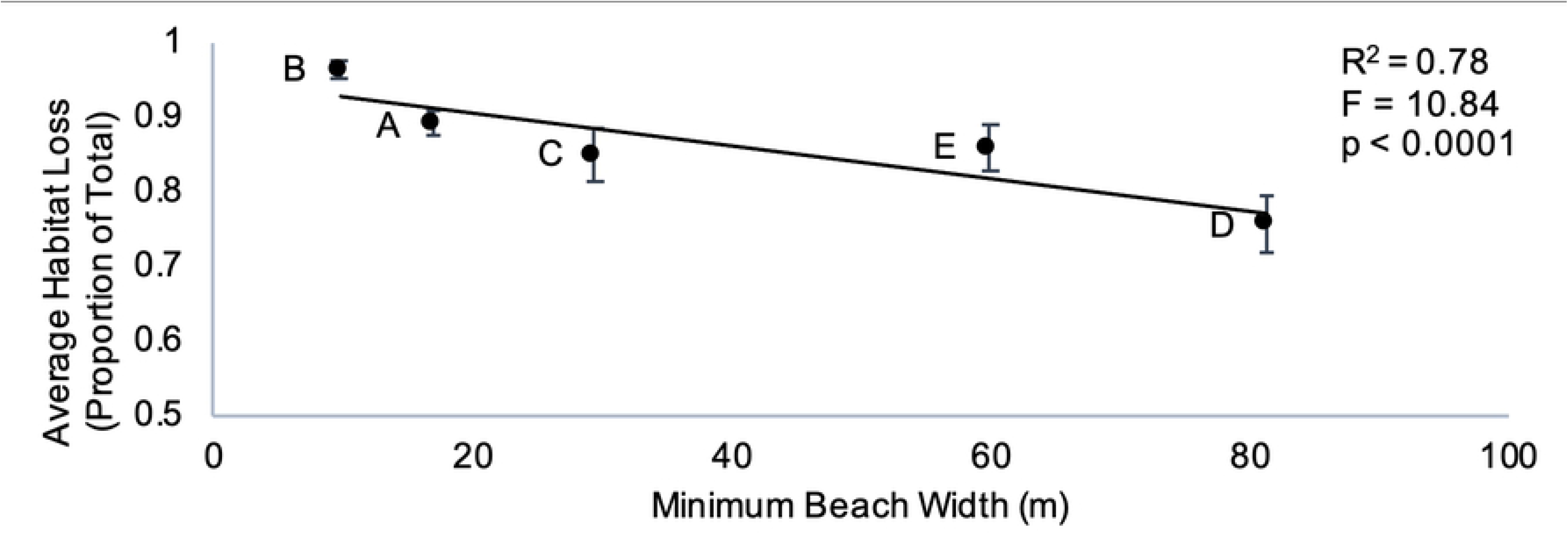
Beach Width versus Habitat Loss. Relationship between minimum beach width (m) and average habitat loss (proportion of whole). Error bars are standard error from the mean.

**Fig 5.**
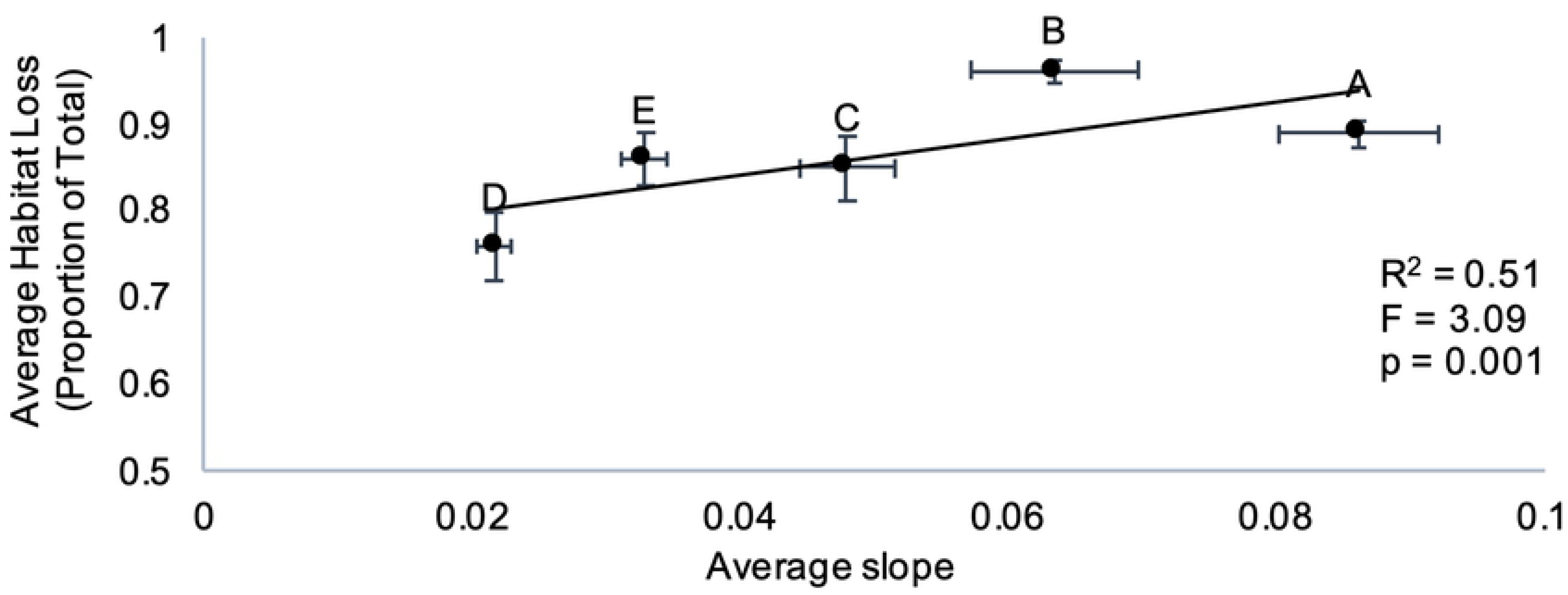
Beach Slope versus Habitat Loss. Relationship between the average slope on beaches A, B, C, D, and E and their average expected habitat loss (expressed in proportion of whole nesting habitat currently available). The averages are for scenarios 0.4, 0.47, 0.48, and 0.63 m for years 2081-2100. Error bars are standard error from the mean.

**Fig 6.**
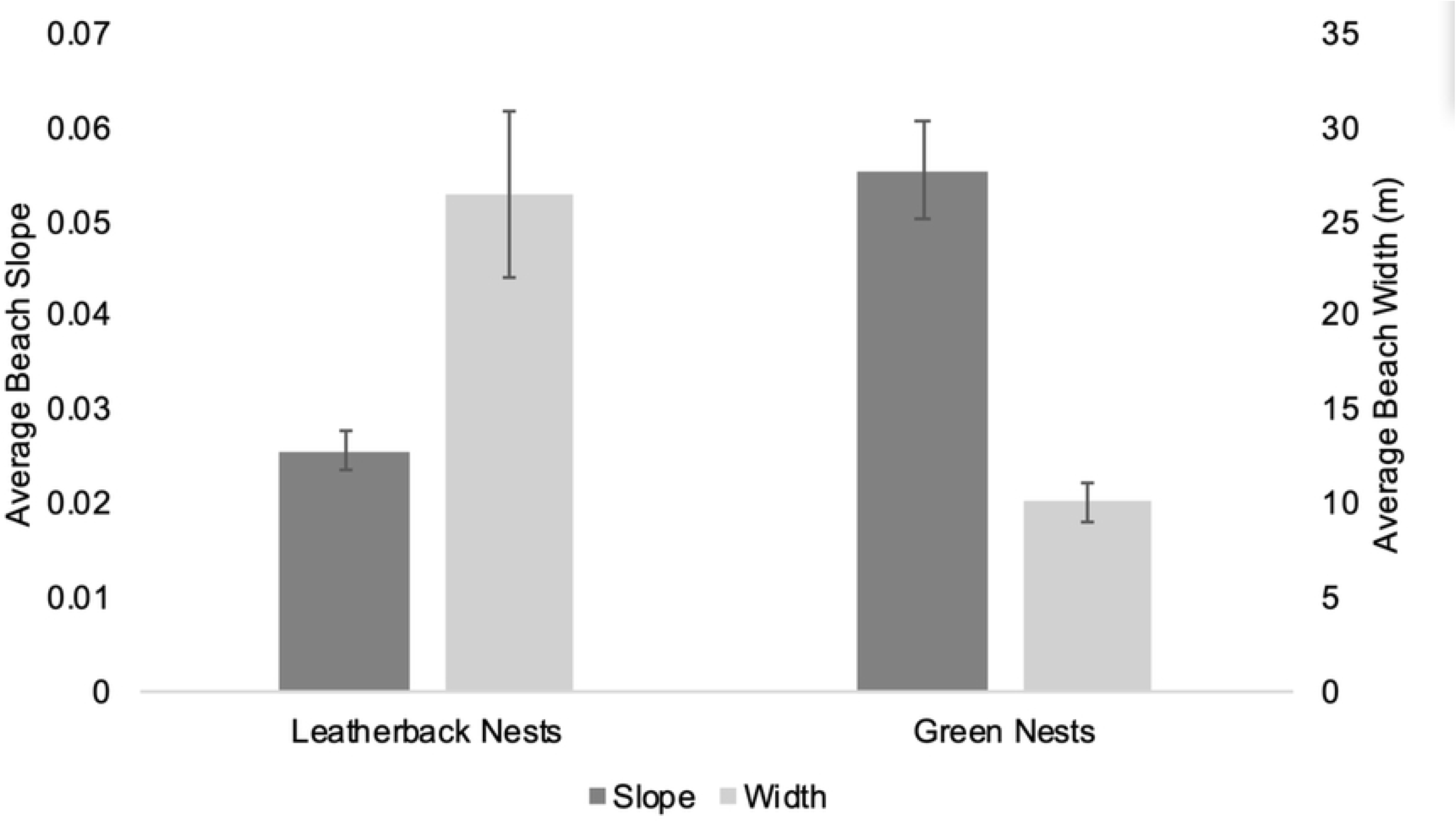
Species Specific Nest Site Selection. Mean beach slope and width for 24 leatherback and 11 green nests on Beaches C & D. Error bars are standard error from the mean.

Threats of nest inundation, predation, and entanglement were identified and uncharacteristic green turtle nesting at the high tide line and in front of the vegetation line in the presence of vegetation berms was documented. With increasing frequency, green sea turtles have been observed entangled in overlying root matrices where the beach has eroded away underneath. Furthermore, a leatherback turtle was discovered with a tree stuck in between her shoulder and neck, causing immobilization. With increased beach erosion and the loss of beach zoning between the high tide line and vegetation line, turtles emerge from the surf in search of dry sand to lay their eggs and instead enter the forest. The analyses predicted that further mortality and hindrances to reproductive success due to these threats are inevitable. Data from standard monitoring efforts during the 2016/17 nesting season showed 89% (n=26) of the time when a leatherback was found digging her nest below the HTL and the nest filled with water, she aborted that nest to choose a drier location closer to the vegetation.

## Discussion

As the beach erodes, beach morphological changes produce abnormal threats to both species of sea turtles as they search for dry sand to lay their eggs. Green turtle nesting beaches will likely undergo more inundation and more rapid erosion due to their narrower, steeper characteristics. With rapid beach erosion on narrow beaches, steep vegetation berms, where the high tide and vegetation lines are one in the same, are left as evidence that the rate of shoreline retreat lags behind that of beach erosion. Green turtles often struggle to reach the vegetation line, as they are unable to surmount steep vegetation berms or become entangled in overhanging root systems where the sand has eroded away beneath (Beach A). Green sea turtles are being found in dangling root matrices with increasing frequency (Honarvar, *personal observations).* Instead of surmounting vegetation berms, green turtles on Beaches A and B have been observed nesting in front of the vegetation and along the high tide line, where their nests are at an increased threat from tidal inundation. Large-scale nest inundation and increased nest conductivity is being observed for this species on Beaches A and B and is expected to continue [32]. The behavior of green turtles to select for narrower, steeper beaches to nest could put them at a greater risk to climate change than other turtle species, as the beaches where they characteristically nest may be morphologically predisposed to erode first based upon the data presented here. Nesting in front of the vegetation line to avoid stark vegetation berms and increased nest conductivity are quantifiable changes within green nesting habitat and nest selection that require further investigation to determine their effects on hatching success and hatchling production. At this time, we expect that increased inundation risk will result in increased nest mortality, and increased sand conductivity will be a significant negative influence on hatching success [32].

In other areas scattered along Bioko’s nesting beaches, classic beach zoning between the high tide line and vegetation is nonexistent but no berms are present, causing the waves to lap against the trees. In these flatter areas, more characteristic of leatherback nesting beaches, like Beaches D and E, leatherback turtles seeking dry sand to lay their eggs can be found stuck in between the trees. Leatherback turtle nest site selection behavior, in nesting closer to the HTL and in front of the vegetation line, puts the nests of this species at immediate risk to tidal inundation as the sea rises [32].

The results presented here represent passive flooding scenarios and the threat of coastal squeeze, which occurs when beaches are obstructed from natural landward movement with increased SLR. Predictions for shoreline retreat, like the Bruun Rule, are controversial and often overly simplified [33]. Modeling shoreline retreat using a Bayesian network has been fruitful in prior studies [34]. Along with shoreline retreat, other factors will undoubtedly play a role in shaping these beaches in the future, such as the effects of long-shore drift and the corresponding reallocation of sediments across nesting beaches, wave heights, the potential net loss of offshore sediment, offshore substrate structure, ocean currents, and increased deposition of sediment materials onto current beach habitat during high tidal inundation events [19]. Complex coastal dynamic processes can be expected to alter the morphology of these beaches with some level of shoreline retreat, but these intricate processes have yet to be studied on Bioko and thus render more complex SLR predictive methods incompatible at this time. The ability of surrounding coral reef growth to correlate with increasing sea levels will likely play a key role in the level of sand accretion seen in the upcoming century [19]. IPCC RCP SLR scenarios for the mid to late 21^st^ century are relative to the reference period of 1986-2005, meaning that the results displayed here could be an estimate applicable for at least 11 years after the official year ranges for reported projections [8]. Inconsistencies in total beach area can be attributed to rounding of proportions and slight changes in model resolution. These predictions can be viewed as the best available insight into the future effects of SLR on Bioko’s five nesting beaches.

The presence of beach sections on Beaches A and B where nesting habitat between the high tide line and vegetation line has already been completely lost is evidence that even though shoreline retreat will occur over time, the rate of beach erosion is currently faster than the rate of shoreline retreat. Unsurmountable vegetation berms have been left as verification of the discrepancy between beach erosion processes and shoreline retreat. There are no anthropogenic barriers to the landward movement of Bioko’s beaches, but rock walls (Beach A) and rivers (Beaches C and D) could be natural barriers [1, 2]. It is likely that a section of Beach A at least 650 m in length will eventually disappear altogether with no inland retreat due to a large rock face directly adjacent to the beach. The face is located farther and farther inland as one moves Southeast along the beach, is at least 50 m tall, and can already be considered the “vegetation line” in some areas.

With increasing SLR, increases in the amount of erosion and flooded nests are expected [11] and have been observed already on Bioko. With an overall reduction in nesting habitat, the density of nests will increase within the area of available nesting habitat. If rains increase in the West African region with climate change, as is suggested with low to medium confidence [35], these beaches are particularly susceptible to increased inundation risk due to rising water tables from both landward and seaward sides. Specific predictions include an increase in the quantity of days experiencing extreme rainfall in West Africa and increased frequency and intensity of rainfall events in the Guinea Highlands and Cameroon Mountains [36–39]. Bioko Island, one of the wettest places in Africa [40], is made up of a complex network of rivers, waterfalls, and lagoons that intersect the beaches at countless points along the shoreline. The fate of one inundated leatherback nest on Beach D is attributable to the high-water table, resulting from a waterfall located directly behind this particular portion of the beach [32].

Creating a hatchery could be an important conservation measure undertaken to protect nests that are likely to be saturated by the tides or rising water tables. Hatcheries have increased the hatching success of sea turtle species on beaches where various anthropogenic and natural threats have made successful in-situ incubation unlikely [41–44]. Although translocating nests can negatively affect embryo development, the relocation of otherwise doomed eggs to a hatchery can result in a net gain in hatchlings produced over time [45].

Increased nest inundation will undoubtedly cause decreases in the reproductive output of all sea turtle species [19]. Preliminary predictions that can be made about species-specific vulnerability with increasing SLR are imperative in understanding which species are at greater impending risks with continued climate change. The data suggest that Beach D will be the beach to maintain the largest amount of nesting habitat for the longest period of time, making it theoretically the most vital beach to protect on the entire island. Unfortunately, it is also the beach that is most threatened by the road built in 2014 and corresponding increase in construction planning, tourists, and illegal egg and adult turtle take (Honarvar et al. 2016). Recommendations have been made to the government of Equatorial Guinea to protect Bioko’s nesting beaches, and especially Beach D, by minimizing development in the Grand Caldera and Southern Highlands Scientific Reserve and the southern beaches, investing in increased tourist environmental awareness campaigns, and increasing enforcement of existing regulations. With minimal development, natural shoreline retreat will have a chance to preserve intricate beach zoning as the sea level rises. By reporting our findings that the beach that is the least vulnerable to future increases in sea level is also the most vulnerable to anthropogenic encroachment, Beach D, the government of Equatorial Guinea can make more informed decisions about the protection of their endangered wildlife.

It was observed that sea turtles have the potential to adapt and choose locations further up the beach, as 89% (n=26) of the time when a leatherback was found digging her nest below the HTL and the nest filled with water, she aborted that nest to choose a drier location closer to the vegetation. This behavior exhibits basic adaptability capabilities. As turtle species can shift their nesting grounds when faced with unsuitable nesting habitats [18], it is important to investigate multiple nesting areas within marine turtle subpopulations [3]. It is possible that turtles could begin to nest on Beach D more frequently, as the other surrounding beaches experience more nesting habitat loss. Leatherback sea turtles may also utilize other nesting habitats on mainland Equatorial Guinea or Gabon more regularly, depending on the extent of beach erosion.

To our knowledge this is the first study to predict the impacts of SLR on sea turtle nesting habitat in Africa and one of the first for a critically important leatherback nesting aggregation worldwide. In the future, advances in modeling methods and increased knowledge of complex coastal processes could be used to improve presented estimates of SLR. These present findings provide a baseline for continued coastal change and habitat use modeling. This study will call attention to the fragility of sea turtle nesting habitat globally and the findings will be valuable to the government of Equatorial Guinea in future developmental planning.

## Acknowledgments

We would like to thank the government of Equatorial Guinea, Universidad Nacional de Guinea Ecuatorial, Instituto Nacional de Desarrollo Forestal y Gestión del Sistema de Áreas Protegidas (INDEFOR-AP), Gertrudis Ribado Mene, The Leatherback Trust and our national and international field assistants (Francisco Ekang Mba Abaga, Juan Jose Edu Abeso Ada, Kenny Ambrose, Ruth Bower-Sword, Brian Dennis, Emily Mettler, Pergentino Ela Nsogo Oye, Adam Quade, Jonah Reenders, Lindsey Rush and Alexis Weaver) for their help and support.

